# Nucleosome binding relinquishes the association of the BAH domain of Orc1 with Sir1

**DOI:** 10.1101/2025.03.14.643217

**Authors:** Heli Jiang, Cong Yu, Chao-Pei Liu, Xiaonan Han, Jingjing Chen, Zhenyu Yu, Rui-Ming Xu

## Abstract

Mating-type switching in *S. cerevisiae* requires silencing of the homothallic mating (HM) loci through formation of position-dependent, gene-independent repressive chromatin domains, resembling heterochromatic regions in higher eukaryotes. Genetic and biochemical studies have identified *cis*-acting DNA elements, called silencers, and *trans*-acting protein factors important for the establishment and maintenance of the silent chromatin. Yet, the molecular mechanism governing the position-dependence of gene silencing is not fully understood. Here we report that the BAH domain of Orc1, which is responsible for recruiting Sir1 to the Orc1-bound silencers, ceases to bind Sir1 in the presence of nucleosome. This finding suggests a unified role of sensing the chromatin environment by Orc1’s BAH domain in transcriptional silencing and specification of replication origins. We further dissected the structural determinants of the BAH domain required for binding Sir1. These results expanded the understanding of Orc1’s functions in epigenetic silencing of the HM loci.

## Introduction

Epigenetic inheritance involves establishment and maintenance of higher-order, repressive chromatin domains. Genetic and biochemical studies of transcriptional silencing of the cryptic mating-type loci, *HML*α and *HMR*a, in *S. cerevisiae* have provided valuable insights into the highly-regulated processes (***Grunstein and Gasser, 2013; Rusche et al., 2003***). Establishment of transcriptional silencing entails the recruitment of silencing proteins to the target genomic loci. In *S. cerevisiae*, both the *HML* and *HMR* loci are flanked by *cis*-acting DNA elements known as the silencers, which contain binding sites for sequence-specific DNA binding proteins, including the origin recognition complex (ORC), Rap1 and Abf1. In turn, these proteins interact with Silence Information Regulator (Sir) proteins, and the recruited Sir proteins spread from the silencers, resulting in the establishment of repressed chromatin domains in the genomic regions flanked by the silencers (***Gartenberg and Smith, 2016; Grunstein and Gasser, 2013; Rusche et al., 2003***). Orc1, the largest subunit of ORC, is responsible for Sir1 recruitment via direct interaction through its Bromo Adjacent Homology (BAH) domain (***Bell et al., 1995; Triolo and Sternglanz, 1996***). Other subunits of the ORC complex also contribute to the recruitment of Sir proteins, but Orc1 appears to function through a distinct pathway exclusively through its BAH domain, as an ectopically tethered Sir1 can bypass the requirement of ORC and initiates silencing (***Bell et al., 1993; Chien et al., 1993; Foss et al., 1993; Fox et al., 1997; Fox et al., 1995; Micklem et al., 1993***). Consistent with the importance of the Orc1-Sir1 interaction, removal of the BAH domain of Orc1 (Orc1^BAH^) displays silencing defects resembling that of *sir1*Δ (***Zhang et al., 2002***).

In addition to its role in binding Sir1, Orc1^BAH^ has also been shown to bind the nucleosome (***Onishi et al., 2007***). The nucleosome binding property of Orc1^BAH^ is shared by the highly homologous BAH domain of Sir3 (Sir3^BAH^) (***Connelly et al., 2006; Onishi et al., 2007***). Their nucleosome binding abilities are greatly enhanced by acetylation of their N-termini, which are catalyzed by the Ard1-Nat1 N-terminal acetyltransferases (***Sampath et al., 2009***). While nucleosome binding by Sir3^BAH^ is well perceived for its function of spreading along the silenced chromatin domain, the nucleosome binding property of Orc1^BAH^ is poorly understood in the context of transcriptional silencing (***Buchberger et al., 2008; Connelly et al., 2006***). In comparison, the association of Orc1^BAH^ with nucleosome is better appreciated in the context of chromosomal replication. The BAH domain was shown to be necessary for stable association of ORC with yeast chromosomes, and important for the activity of a class of origins with a distinct local nucleosome structure (***Muller et al., 2010***). This observation is also mirrored in the finding that certain chromosomal features in addition to the autonomously-replicating consensus sequences (ACS) are important for ORC binding and origin function (***Eaton et al., 2010***). It is interesting to wonder if the nucleosome binding property of Orc1^BAH^ also contributes to the recognition of specific chromatin features surrounding the silencers that specify the position dependence of transcriptional silencing.

A related, yet incompletely understood matter concerning Orc1^BAH^ is its critical determinants for Sir1 binding. It was shown previously that a segment called the H-domain within the BAH domain was necessary for Sir1 binding. Replacing the H-domain of Sir3^BAH^ with that of Orc1^BAH^ in the context of full-length Sir3 enables it to bind Sir1 (***Zhang et al., 2002***). However, this binding has not been demonstrated for the BAH domain alone *in vitro*. Now the crystal structure of Orc1^BAH^ in complex with the C-terminal Orc1-interacting domain (OID) of Sir1 is available, it should facilitate a better understanding of the structural determinants for Sir1 binding (***Hou et al., 2005; Hsu et al., 2005***).

Here we show that nucleosome impedes Sir1 binding by Orc1^BAH^, indicating the need of a suitable chromatin environment for Sir1 recruitment in yeast cells. We also uncovered an amino acid difference between Orc1^BAH^ and Sir3^BAH^ that is crucial for their Sir1 binding properties. Altogether, these findings should facilitate detailed mechanistic understandings of transcriptional silencing of the mating-type genes in *S. cerevisiae*.

## Results

### Nucleosome disrupts Orc1-Sir1 interaction

Orc1^BAH^ was previously shown to interact with the nucleosome core particle (NCP) in a manner dependent on acetylation of its N-terminus. Several structures of the homologous Sir3^BAH^ in complex with NCP have been solved, and the structures revealed that the BAH domain makes extensive contact with all four histones exposed on one side of the disk-shaped nucleosome surface (***Armache et al., 2011; Arnaudo et al., 2013; Wang et al., 2013; Yang et al., 2013***). Besides the histone core domains, a segment of the N-terminal tail of histone H4, spanning residues ∼15-23, engages Sir3^BAH^ and stabilizes the Sir3-NCP complex. For the crystal structure of the Orc1^BAH^-NCP complex, Orc1^BAH^ binds NCP very similar to that of Sir3^BAH^, despite three notable differences (***De Ioannes et al., 2019***). First, the ordered N-terminal tail of histone H4 starts at residue 18, consistent with the report that the Orc1^BAH^ binding to NCP is insensitive to H4K16 acetylation; Second, the loop binding the acidic patch of NCP is only partially ordered in Orc1^BAH^; Third, the loop connecting the two helices of the Sir1-binding H-domain is disordered.

Since Sir1^OID^ forms a complex with Orc1^BAH^, we wondered if the BAH domain would choreograph the binding to Sir1^OID^ and NCP. Docking of the Orc1^BAH^-Sir1^OID^ crystal structure onto NCP through superposition of the BAH domain revealed that a ternary Orc1^BAH^-Sir1^OID^-NCP complex is permissible (Figure 1A)(***De Ioannes et al., 2019; Hou et al., 2005; Hsu et al., 2005***). To validate it experimentally, we first prepared the Orc1^BAH^-Sir1^OID^ complex by mixing insect cell-expressed Orc1^BAH^, which is N-terminal acetylated and binds NCP on its own, with bacterially expressed Sir1^OID^. The binary complex was then mixed with reconstituted NCP and applied to a gel-filtration column to isolate the putative ternary complex. Surprisingly, even at 50 mM NaCl concentration, the Orc1^BAH^-Sir1^OID^ complex did not co-elute with NCP (Figures 1B & 1C). Instead, a portion of Orc1^BAH^ co-eluted with NCP. It is not clear whether the Orc1^BAH^ co-eluted with NCP was due to an excess or was disrupted from the Orc1^BAH^-Sir1^OID^ complex by NCP. To discern the possibilities, GST-tagged Sir1^OID^ was immobilized on glutathione sepharose resins and first incubated with purified Orc1^BAH^. Subsequent addition of NCP shows that the amount of bound Orc1^BAH^ is significantly reduced (Figure 1D), indicating that NCP snatches Orc1^BAH^ away from GST-Sir1^OID^. Furthermore, excessive amount of Sir1^OID^ added to the pre-formed NCP-Orc1^BAH^ complex cannot compete with NCP for Orc1^BAH^ (Figure S1), even though an Orc1^BAH^-Sir1^OID^ complex is more resilient to higher salt concentration than the NCP-Orc1^BAH^ complex does (***Hou et al., 2005; Hsu et al., 2005***). These observations suggest that distinct regions of Orc1^BAH^ are involved in binding Sir1^OID^ and NCP, otherwise the more stable Orc1^BAH^-Sir1^OID^ complex would prevail.

**Figure 1.**
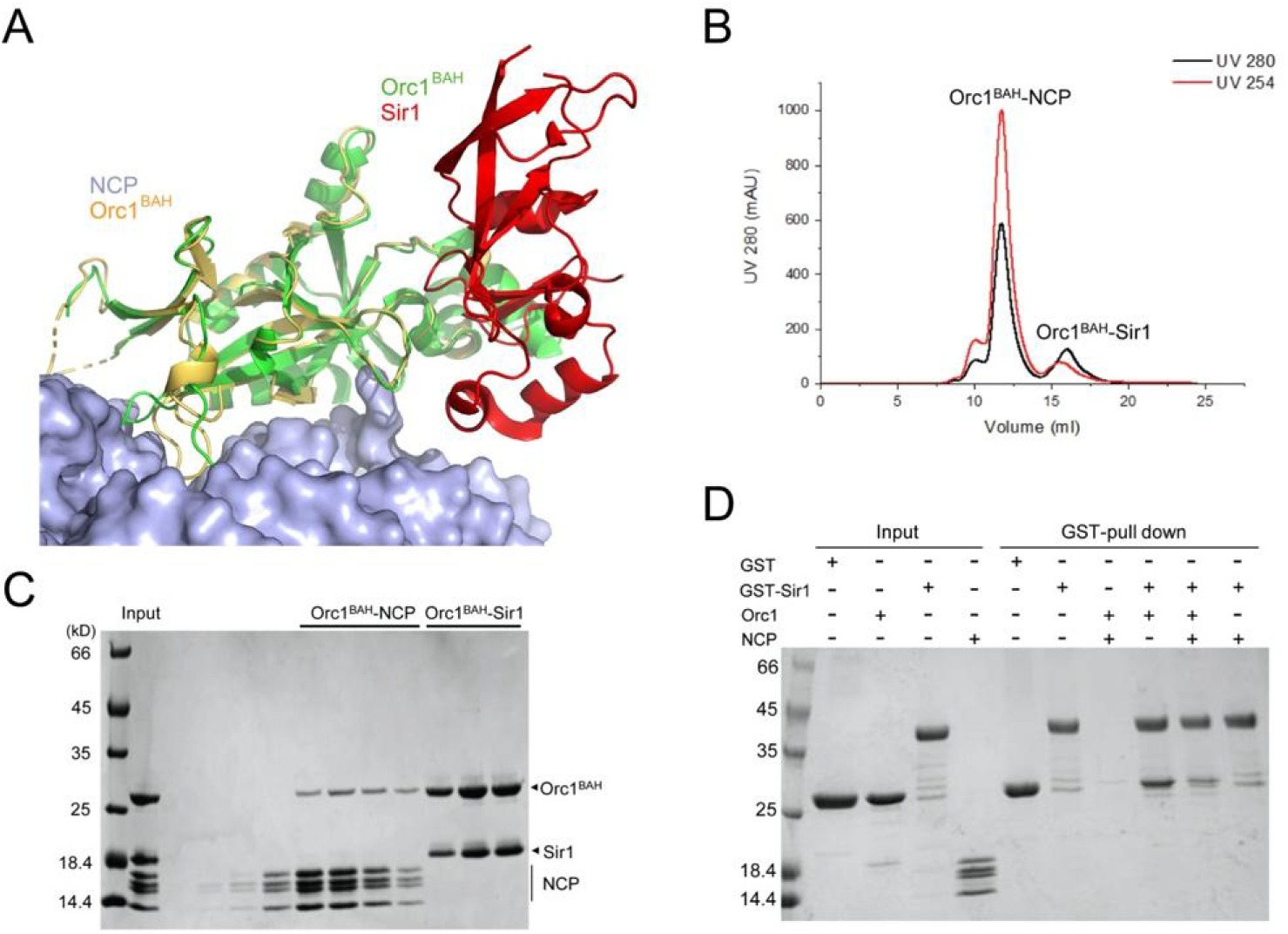
Nucleosome disrupts the binding of Orc1 to Sir1. (A) Superposition of the structures of Orc1^BAH^-Sir1^OID^, shown as green and red ribbons, respectively, and the NCP-Orc1^BAH^, shown in blue surface and yellow ribbon representations, demonstrate the feasibility of an NCP-Orc1^BAH^-Sir1^OID^ complex. (B) UV absorbance chromatogram, with absorbances of 280 nm and 254 nm shown in black and red lines, respectively, of a size-exclusion column run shows that Sir1 does not co-elute with NCP. (C) SDS-PAGE gel shows the eluted fractions from the size-exclusion column. (D) GST pulldown assay shows that NCP disrupts the Orc1^BAH^-Sir1^OID^ interaction.

### Cryo-EM structure of the Orc1^BAH^-NCP complex

It is puzzling that the Orc1^BAH^-NCP and Orc1^BAH^-Sir1^OID^ structures permit the formation of an Orc1^BAH^-Sir1^OID^-NCP complex, yet our biochemical results contradict the prediction. To exclude the possibility that conformational bias caused by crystal packing misleads the prediction, we carefully examined the crystal packing contacts in both the Orc1^BAH^-Sir1^OID^ and Orc1^BAH^-NCP structures. The Orc1^BAH^-Sir1^OID^ structure was solved from crystals belonging to two different spacegroups, and the intermolecular contacts across the crystallographic asymmetric units are different (***Hou et al., 2005; Hsu et al., 2005***). Yet, the Orc1^BAH^-Sir1^OID^ interfaces are remarkably similar in the two structures, indicating little perturbation by crystal packings. In the Orc1^BAH^-NCP crystal structure, a major Sir1-interacting region in the Orc1 BAH domain, called the H-domain, interacts with nucleosomal DNA of neighboring NCPs within and across the asymmetric unit (Figure S2) (***De Ioannes et al., 2019***). To evaluate the potential impact of crystal packing, we determined the Orc1^BAH^-NCP structure by single-particle cryo-electron microscopy (cryo-EM). Complexes of Orc1^BAH^ bound to NCP at 1:1 and 2:1 molar ratios were observed, and their structures were determined at 3.5 Å and 4.0 Å resolutions, respectively (Figures 2A, S3, S4 and Table S1). In both structures, the NCP densities are well defined, and that of the BAH domains are weaker. In particular, the densities for the Sir1-interacting H-domain and several peripheral loops of Orc1^BAH^ are poorly defined. Improvement of the EM density with refinement of the masked volumes enclosing the BAH domain allowed unambiguous docking of Orc1^BAH^ with NCP.

**Figure 2.**
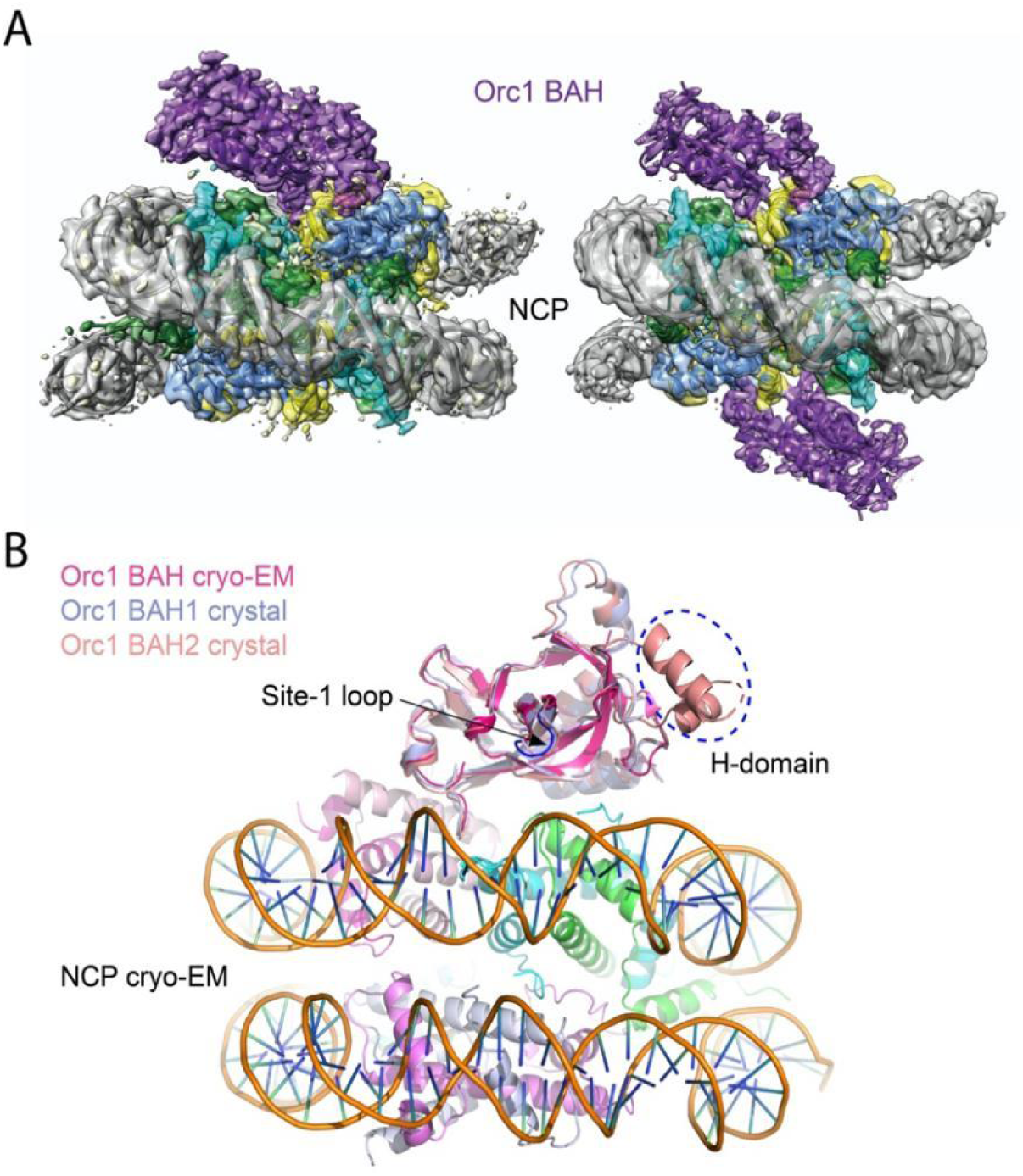
Structure of Orc1^BAH^ bound to NCP. (A) Left panel, 3.5 Å cryo-EM structure of one Orc1^BAH^ molecule bound to one side of NCP; Right panel, 4.0 Å cryo-EM structure with one Orc1^BAH^ bound to each side of NCP. The densities colored in purple are that of Orc1^BAH^, and those of DNA and histones are colored in grey and miscellaneous colors, respectively. (B) The density for Orc1^BAH^ (magenta) is generally weaker than that for NCP, and the H-domain of Orc1^BAH^ is not visualized. Two Orc1^BAH^ molecules bound to the same NCP in the crystal structure (PDB ID: 6om3) are superimposed. One of the Orc1^BAH^ molecule from the crystal structure, colored wheat, contains a partially ordered H-domain, while the other, colored light blue, has a disordered H-domain, as in the cryo-EM structure. The H-domain and the Site-1 loop, two important regions for Sir1 binding, are indicated with a dashed-line oval and an arrow head, respectively.

Nevertheless, the density for the H-domain of Orc1 still cannot be modeled, and the same is true in the crystal structure, as only one of the two Orc1^BAH^ molecules bound to each NCP contains partially ordered H-domain, indicating dynamic nature of this region in solution (Figures 2B & S2). Given the flexibility of the H-domain in Orc1^BAH^, there appears to have no a priori reason for Sir1 not to bind Orc1^BAH^ and forms a ternary complex with NCP. The discrepancy between the structural prediction and the biochemical data calls for careful reexamination of critical elements required for Orc1^BAH^-Sir1^OID^ interaction.

### BAH domain elements important for interaction with Sir1

The main Sir1-contacting area is located in the H-domain of Orc1^BAH^, wherein lies two of the three Sir1^OID^-interacting regions, denoted sites II and III (***Zhang et al., 2002***). The H-domain residues directly involved in Sir1 binding include Arg92, Trp93, Phe94, Asn117, Asn120, Lys121, Ser124 and Glu125 (Figure 3A). However, the exact identity of many of the residues is not required for interaction with Sir1, as they are also present in the non-Sir1 binding Sir3 BAH domain. The most different residue is Ser124 in Orc1^BAH^ and Phe124 in Sir3^BAH^. But Ser124 of Orc1^BAH^ faces a hydrophobic pocket formed by Tyr489, Leu504 and Phe557 of Sir1, substituting it with a phenylalanine should not compromise the Orc1^BAH^-Sir1^OID^ interaction (Figure 3A). Indeed, the S124F mutant of Orc1^BAH^ can be pulldown by GST-Sir1^OID^ as efficiently as the wild-type Orc1^BAH^ (Figures 3B, lanes 3 & 5; Figure S5).

**Figure 3.**
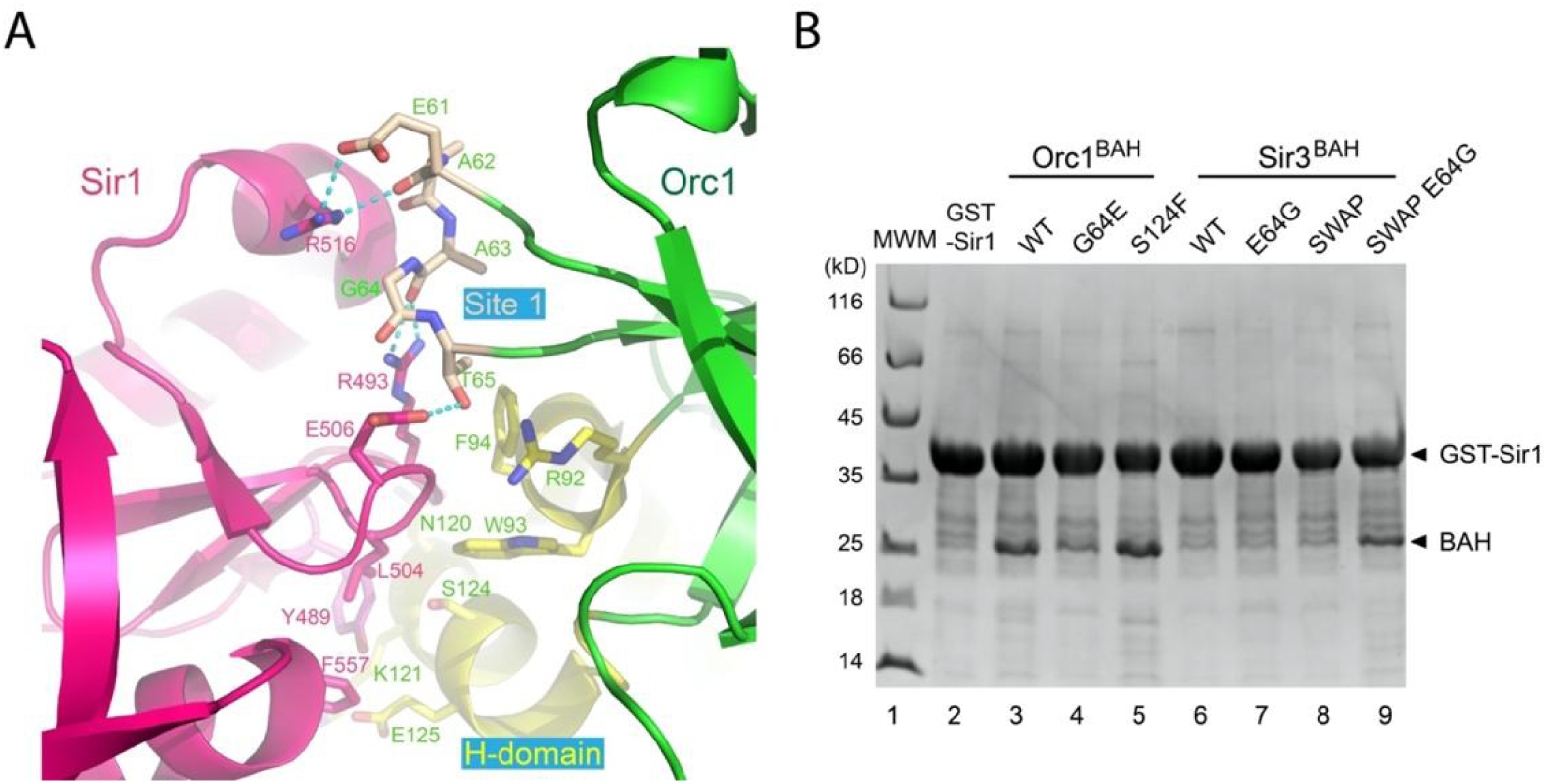
Key elements for BAH domain-Sir1 interaction. (A) Depiction of Orc1^BAH^-Sir1^OID^ interaction (PDB ID: 1zbx), with Orc1 and Sir1 colored in green and red, and the H-domain and Site 1 regions of Orc1^BAH^ are highlighted yellow and beige, respectively. Key residues involved in their interactions are shown in a stick model and labeled. (B) GST-Sir1 pulldown of indicated Orc1^BAH^ and Sir3^BAH^ variants produced by co-expression shows the importance of both the H-domain and Site 1 residues in Sir1 binding. The inputs for the pulldowns are shown in Figure S5.

We have previously demonstrated that replacing the entire H-domain of the full-length Sir3 with that of Orc1^BAH^ enables the chimeric Sir3 to bind full-length Sir1 in vitro (***Zhang et al., 2002***). However, the presence of extra protein domains in full-length Sir3 and Sir1 complicates the interpretation of the binding effect of the H-domain (***Connelly et al., 2006; Hou et al., 2009***). Here we transplanted the entire H-domain of Orc1^BAH^ to Sir3^BAH^, yielding a chimeric protein, termed Sir3^BAH^-swap, with residues 96-130 of Sir3^BAH^ replaced by the corresponding Orc1^BAH^ fragment. However, Sir3^BAH^-swap cannot be pulldown by GST-Sir1^OID^ (Figure 3B, lane 8), indicating that additional critical elements located outside of the H-domain of Orc1^BAH^ is also needed for efficient Sir1 binding.

The only Sir1-contacting region outside the H-domain in Orc1^BAH^ is Site I, which is located at the tip of the loop connecting β4 and β5 of the BAH domain core (Figure 3A). Its Glu61 and Ala63 interact with two Sir1 arginines Arg516 and Arg493, respectively, via mainchain carbonyl groups, while the sidechain carboxylate group of Glu61 also makes a hydrogen bond with the guanidino group of Arg516. The hydroxyl group of Thr65 makes a hydrogen bond with the carboxylate group of Sir1 Glu506, which, however, is conserved in Sir3. Ala63 is neighbored by Gly64, but the corresponding residue in Sir3 is Glu64. Surprisingly, a G64E mutation of Orc1^BAH^ noticeably weakened the interaction with Sir1 (Figure 3B, lane 4). A reverse mutant of Sir3, E64G, does not show obvious binding to Sir1 (lane 7). However, a combination of E64G and Sir3^BAH^-swap mutations gains the ability to interact with Sir1 (Figure 3B, lane 9). These analyses indicate that both the H-domain and the conformation of the β4-β5 loop are crucial for interaction with Sir1.

### Nucleosome-regulated interaction between Orc1 and Sir1

The above analysis revealed that coordinated interaction with the H-domain and the β 4-β5 loop of Orc1^BAH^ is essential for stable binding of Sir1^OID^. In the structure of the Orc1^BAH^-NCP complex, the H-domain is mostly disordered, indicating a dynamic but accessible protein region for Sir1^OID^ binding, while the β4-β5 loop is well ordered. In fact, this loop is located at the end of the conserved histone H4 tail binding cleft, which is formed between NCP and Orc1^BAH^ or Sir3^BAH^. In the Orc1^BAH^-NCP complex, the first ordered N-terminal residue of histone H4 is His18, while it starts at Ala15 in the Sir3^BAH^-NCP complex (Figure 4A). Ala15 of histone H4 is within 4 Å from the β4-β5 loop of Sir3^BAH^, and based on the similar binding trajectory of the histone H4 tail, the N-terminal disordered portion is expected to be in the general vicinity of the β 4-β 5 loop of Orc1^BAH^. Could it be that the dynamic N-terminal portion of the histone H4 tail interfere with Sir1^OID^ binding? To test this possibility, we reconstituted NCPs with H4 variants having 15 or 20 N-terminal residues removed and tested their binding to Orc1^BAH^ and Sir1^OID^. We find that the binding between Orc1^BAH^ and NCP is maintained with N-terminal 15 residues of H4 removed, but Sir1^OID^ still cannot form a ternary complex with Orc1^BAH^ and the NCP (Figure 4B). Orc1^BAH^ itself no longer binds the NCP with 20 N-terminal residues of histone H4 removed (Figure S6). These findings point to the cause that the movement of the β4-β5 loop away from the Sir1 arginine residues by ∼ 1 Å due to NCP binding creates a setting that the β4-β5 loop and the H-domain cannot coordinate the binding of Sir1^OID^ (Figure 4A).

**Figure 4.**
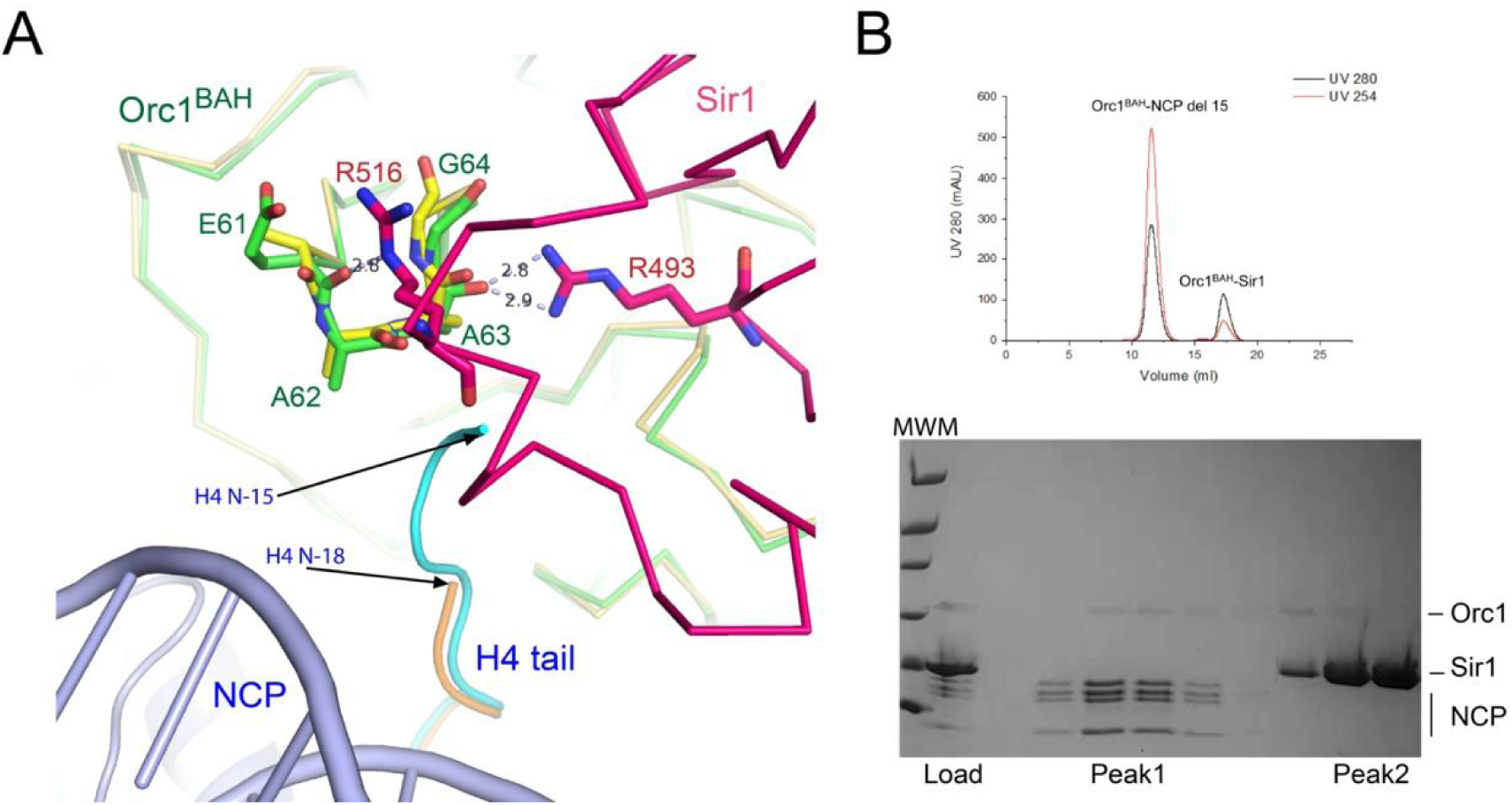
Molecular basis for NCP regulated Orc1^BAH^-Sir1^OID^ interaction. (A) The N-terminal tail of histone H4 reaches near the Sir1-binding Site 1 of Orc1^BAH^. The H4 tail shown in orange is from the Orc1^BAH^-NCP complex (PDB ID: 6om3), and it is visible starting from residue-18 at the N-terminus. The H4 tail from residue-15 of the Sir3^BAH^-NCP complex (cyan) is superimposed. Alignment of Orc1^BAH^ structures from its complexes with NCP and Sir1, shown in yellow and green, respectively, reveals displacement of the carbonyl groups of Glu61 and Ala63. Sir1^OID^ model from the complex with Orc1^BAH^ is shown in red, and its Arg493 and Arg516 interacting with Ala63 and Glu61 are depicted in a stick model. (B) Size-exclusion column chromatography shows that NCP reconstituted with deletion of 15 N-terminal residues of histone H4 still binds Orc1^BAH^ and resists the formation of a complex with Sir1^OID^.

## Discussion

To date, the only known mechanism by which the BAH domain of Orc1 functions in transcriptional silencing is through its direct interaction with Sir1 (***Bell et al., 1995; Triolo and Sternglanz, 1996; Zhang et al., 2002***). The discovery of the nucleosome binding property of Orc1^BAH^ and Sir3^BAH^ gives rise to the question of whether and how this property may be related to their silencing functions (***Onishi et al., 2007***). This property of Sir3 is well anticipated, as it spreads along the nucleosomes to form the repressive chromatin domain, and the BAH domain in place of the full-length Sir3 can achieve partial silencing (***Buchberger et al., 2008; Sampath et al., 2009***). In comparison, the BAH domain of the silencer-bound Orc1 is restricted to the vicinity of the silencer, and it will most likely only encounter nucleosomes flanking the silencer.

Our study here provides an interesting insight into the nucleosome binding property of Orc1^BAH^ in the context of transcriptional silencing. Our finding implies that the silencer-bound Orc1 will not be able to bind Sir1 if nucleosomes are within reach of its BAH domain. The inability of Orc1^BAH^ to bind Sir1^OID^ in the presence of NCP is not due to overlapping binding sites, nor due to steric hindrance by NCP. Previous structural studies of the Orc1^BAH^-Sir1^OID^ complex revealed that the H-domain moves toward the β 4-β 5 loop, where Gly64 is located, upon Sir1^OID^ binding, indicating that a precise spatial arrangement between these elements is required for stable Sir1^OID^ binding. This requirement may also account the inability of Sir3^BAH^ to bind Sir1^OID^, as only both an H-domain swap and a E64G mutation in the β4-β5 loop enable a gain of interaction. Structure comparison shows that in the presence of NCP, while the H-domain is freely available, the β4-β5 loop is displaced with respect to the core of Orc1^BAH^, rendering a nucleosome regulated interaction with Sir1^OID^.

Our discovery indicates that a functional silencer should not reside in an environment too close to nearby nucleosomes. This property of Orc1^BAH^ is reminiscent of its function in the selection of origins of replication (***Muller et al., 2010***). It was originally thought that the BAH domain of Orc1 does no function in DNA replication, but later studies showed that it was necessary for stable association of the ORC complex with chromosomes. Furthermore, the BAH domain sense the local nucleosome structure and distinguishes a class of BAH domain sensitive origins. The ORC binding sites have a characteristic asymmetric pattern of positioned nucleosomes, and the position of the −1 nucleosome affect the origin activity (***Eaton et al., 2010; Muller et al., 2010***). A salient feature of transcriptional silencing is position dependent and gene-independent. We propose that the BAH domain of Orc1 specifies the position dependent property, with nucleosomes flanking the silencers determining the Sir1-dependent silencing activities of these silencers. Nucleosome positions of the HML and HMR loci have been precisely mapped (***Ravindra et al., 1999; Weiss and Simpson, 1998***). Further in-depth genetic and genomic studies should develop detailed mechanistic insights into the silencer-associated function of the Orc1 BAH domain.

## Materials and Methods

### Protein expression and purification

*S. cerevisiae* Orc1 BAH domain (Orc1^BAH^, amino acids 1–219) was expressed as a C-terminal 6xHis-tagged fusion protein in Sf21 insect cells with the pFastBac1 Bac-to-Bac baculovirus expression system (Invitrogen). 72 h after virus infection at 27 °C, cells were harvested by centrifugation and resuspended in the lysis buffer containing 20 mM Tris-HCl, pH 7.5, 300 mM NaCl, and 5% glycerol. Cells were ruptured with sonication and cellular debris were removed by centrifugation. The cleared lysate was then added to Ni-NTA (GE Healthcare) beads pre-equilibrated with the lysis buffer for binding. After successive washing steps with the buffer containing 10, 20 and 50 mM imidazole, the bound proteins were eluted with the same buffer containing 300 mM imidazole. The eluted proteins were desalted, concentrated and further purified on a Superdex 200 10/300 GL size-exclusion column (GE Healthcare) pre-equilibrated with the SEC buffer (20 mM Tris, pH 7.5, 150 mM NaCl, 1 mM DTT, and 1 mM EDTA).

Sir1^OID^ (amino acids 480-612) was expressed as a C-terminal 6xHis-tagged protein in the BL21(DE3) strain of *E. coli* using a pET-23a vector (Novagen), and the protein was purified following the same procedure as that for Orc1^BAH^, except that and the SEC buffer contains 2 M NaCl, 5% glycerol, and no EDTA was added. GST-Sir1^OID^ used for pulldown experiments was produced in *E. coli* using a pGEX-KG vector (Novagen), and purified using glutathione resins, followed by size-exclusion column chromatography in a buffer containing 20 mM Tris at pH 7.5, 200 mM NaCl, 5% glycerol and 1 mM DTT.

For assessment of BAH domain elements important for interaction with Sir1, GST-Sir1^OID^ and Orc1^BAH^, Sir3^BAH^ or their mutants (Orc1^BAH^, Orc1^BAH^G64E, Orc1^BAH^S124F, Sir3^BAH^, Sir3^BAH^E64G, Sir3^BAH^-swap, Sir3^BAH^-swap E64G) were co-expressed using a pGEX-KG and a pMR101 vector, respectively, in the Rosetta (DE3) strain of *E. coli*.

### Preparation of NCP and its complex with Orc1^BAH^

All recombinant yeast histone octamers, except the one with 20 N-terminal residues of histone H4 deleted (H4 del-20), were obtained by co-expression in the BL21(DE3)-RIL strain of *E. coli* (***Kingston et al., 2011***). The octamer with H4 del-20 was prepared by mixing bacterially co-expressed soluble H2A-H2B and refolded H3-H4 del-20, and purified at 2M NaCl through a size-exclusion column. NCP was reconstituted by salt dialysis using the recombinant histone octamers and bacterially produced 147 bp 601 strong positioning DNA sequence following established protocols (***Dyer et al., 2004; Kingston et al., 2011; Lowary and Widom, 1998; Luger et al., 1999***). The Orc1^BAH^-NCP complex was assembled by mixing Orc1^BAH^ and NCP in a 5:1 molar ratio and dialyzed the mixture overnight against 20 mM HEPES, pH 7.5, 50 mM NaCl, 1 mM EDTA and 1 mM DTT, and separated from free NCP and Orc1^BAH^ using a Superdex 200 10/300 GL size-exclusion column (GE Healthcare).

### GST pulldown

First, 10 µg of GST-Sir1^OID^ and Orc1^BAH^-C-His were mixed at a molar ratio of 1:2 in the binding buffer (20 mM Tris, pH 7.5, 50 mM NaCl, 5% glycerol and 1 mM DTT), and incubated for 30 min on ice. Then, 20 µl of glutathione-agarose resins were added and incubated for 2 hours at 4 °C with end-over-end rotation. After washing the resins twice with the binding buffer supplemented with 0.2% NP-40, the protein bound beads were incubated with NCP at two fold molar excess for additional 2 hours at 4 °C with end-over-end rotation. Finally, the resins were washed two more rounds, and the bound components were analyzed by SDS-PAGE. As a control, GST alone, GST-Sir1 alone, the Orc1^BAH^-NCP complex, and the mixture of GST-Sir1^OID^ and NCP were treated in the same way.

For assessment of Sir1^OID^ interaction with Orc1^BAH^ or Sir3^BAH^ variants, GST-Sir1^OID^ was co-expressed with Orc1^BAH^ or Sir3^BAH^ variants in *E. coli* under same conditions. Equal volumes of post-induced bacteria cells were collected and suspended in the binding buffer (20 mM Tris, pH 7.5, 100 mM NaCl, 5% glycerol and 1 mM DTT). After sonication and centrifugation, cellular debris were removed and the cleared lysate were incubated with 20 µl of glutathione-agarose resins in the binding buffer for 2 hours at 4 °C with end-over-end rotation. After washing the resins three times with the binding buffer supplemented with 0.2% NP-40, the bound components were analyzed by SDS-PAGE.

### Cryo-EM specimen preparation and data collection

The purified Orc1^BAH^-NCP complex was first subjected to Grafix crosslinking with glutaraldehyde. Briefly, the protein complex was loaded onto a 5 ml liner 10–30% glycerol gradient in the buffer containing 20 mM HEPES, pH 7.5, 50 mM NaCl, 1 mM EDTA and 1 mM DTT supplemented with 0-0.15% EM-grade glutaraldehyde (Sigma-Aldrich). After centrifugation at 4°C for 16 hr at 38,000 rpm in a SW50 rotor (Beckman Coulter), the sample was manually fractionated into 25 fractions of 200 μL each from top to bottom using pipettes. Crosslinking of each fraction was analyzed on 8% denaturing polyacrylamide gel.

The crosslinked fractions were first examined by negative-staining EM for particle homogeneity. Suitable fractions were changed to a buffer with the glycerol removed for Cryo-EM studies. Cryo-EM specimens were prepared using the 300 mesh Quantifoil R2/1 copper grids. A 3 μL aliquot of the sample at a concentration of ∼1 mg/mL was applied to glow-discharged grids. After incubation for 10 s, excess sample was blotted with filter paper for 6 s and the grid was flash-frozen in liquid ethane using a FEI Vitrobot device. All cryo-EM samples were prepared using the same procedure at 10°C and 100 % relative humidity.

Cryo-EM images were collected on an FEI Talos Arctica electron microscope equipped with a GIF Quantum energy filter and operating at 200 kV with a nominal magnification of 130,000. Images were recorded by a Gatan Bio-Quantum K2 Summit direct electron detector. The slit width for zero loss peak was 20 eV. The camera was in a super-resolution mode with 1 Å physical pixel size (0.5 Å super resolution pixel size). The defocus range was set between −0.8 and −1.2 μm. Each image was exposed for 5 s, resulting in total dose of ∼50 e^-^/Å^2^ and 32 frames per movie stack.

### Cryo-EM image processing

Each movie stack was binned and subjected to motion correction using the MotionCor2 software (***Zheng et al., 2017***). This procedure produced two images for each movie stack by summing all frames with or without dose weighting. The contrast transfer function (CTF) parameters were measured by Gctf (***Zhang, 2016***) using the summed image without dose weighting. Micrographs with poor estimated resolutions were eliminated in the screening. The summed image with dose-weighting was used for further data processing in Relion-3.0 (***Zivanov et al., 2018***).

Dataset-1 contains 2930-micrographs, of which fourteen were randomly selected for auto-picking using the Laplacian-of-Gaussian method in Relion-3.0, generating a small subset containing 12,003 particles. These particles were processed with reference-free 2D classification. Nine 2D class average images were selected as references for automatic particle picking of the complete dataset. A total of 2,345,924 particles were picked from 2,930 micrographs and processed by reference free 2D classification. 1,766,421 particles were selected for further 3D classification with an initial model generated by converting the nucleosome crystal structure (PDB ID:1ID3) to the electron microscopy density using the EMAN2 pdb2mrc.py script (***Tang et al., 2007***) and low-pass filtered to a resolution of 50 Å.

After several rounds of 2D and 3D classifications, 251,525 particles (Class 4A) were selected for further 3D auto-refinement and postprocess with soft mask, which generated a 3D reconstruction of Orc1 BAH-NCP (1:1) complex at an overall resolution of 3.5 Å, estimated based on the criterion of a Fourier shell correlation of 0.143 (***Rosenthal and Henderson, 2003; van Heel and Schatz, 2005***).

Initial reconstruction of the 2:1 Orc1^BAH^-NCP complex was carried out using dataset-2, which contains another 1,627 micrographs. A total of 1,326,260 particles were picked and processed by reference-free 2D classification. After several rounds of 2D and 3D classifications, two higher quality classes were merged and 254,395 particles were selected. Then, the particles are combined with that of dataset-1 used for 3D auto-refinement of the 1:1 complex. Another 3D classification was performed and 119,206 particles were selected for 3D auto-refinement using C1 or C2 symmetry, and the latter yielded a better outcome, with an overall resolution of 4.36 Å. After applying a soft mask and post process, the overall resolution improved to 4.0 Å. The local resolution of the final maps was calculated using ResMap (***Kucukelbir et al., 2014***). A summary of the above procedures and detailed statistics are shown in Figures S3, S4 and Table S1.

### Model building, refinement and validation

Atomic models of the W601 DNA (PDB: 3LZ0), the *S. cerevisiae* histone octamer (PDB: 1ID3) and the *S. cerevisiae* Orc1 BAH domain (PDB: 1M4Z or 6OM3) were docked into the 3.5-Å 1:1 Orc1^BAH^-NCP map using UCSF Chimera (***Goddard et al., 2007***). The model was further adjusted and rebuilt manually in COOT (***Emsley et al., 2010***). The coordinates for the regions where no density was observed were deleted and adjustments were also made according to the actual residues or DNA sequence used in this study. The 4.0-Å 2:1 Orc1^BAH^-NCP model was generated from the 1:1 Orc1^BAH^-NCP model. The models were then real-space refined using PHENIX (***Adams et al., 2010***) with secondary structure restraints. Finally, the models were validated using MolProbity (***Chen et al., 2010***). Refinement statistics are shown in Table S1.

## Accession codes

EM maps and fitted structural models have been deposited in EMDB (EMD-31029 and EMD-31030) and PDB (7E9C and 7E9F).

## Acknowledgements

We thank Dr. Hao-Chi Hsu for early work about mutagenesis of Sir3 and Orc1 BAH domains, Dr. Xinzheng Zhang for advices on cryo-EM structure determination, Drs. Boling Zhu, Xiaojun Huang and Lihong Chen at Center for Biological Imaging (CBI) of Core Facility for Protein Science Research at Institute of Biophysics (IBP), Chinese Academy of Sciences (CAS) for support in cryo-EM data collection. The research was supported by grants from Ministry of Science and Technology of China (2019YFA0508900 and 2018YFE0203300), Natural Science Foundation of China (31991162, 91853204 and 31521002), and the Strategic Priority Research Program of CAS (XDB37010100). Dr. Zhenyu Yu is supported by the Youth Innovation Promotion Association of CAS (2017131).

## Author contributions

HJ, prepared nucleosomes and proteins, performed binding experiments, drafting the article; CY, processed cryo-EM data, built and refined structure models, edited the manuscript; CPL, supervised data acquisition, processing and structure determination; XH, acquisition and processing of cryo-EM data, building and refinement of structure models; JC, assisted experiments and managed the lab; ZY, supervised protein and nucleosome preparation, prepared cryo-EM samples and collected data, analyzed and interpreted data, drafted the article; R-MX, conception and design, analysis and interpretation of data, drafting the article.

## Competing interests

All authors declare no competing interests.

## Supplemental Table and Figures

**Figure S1.**
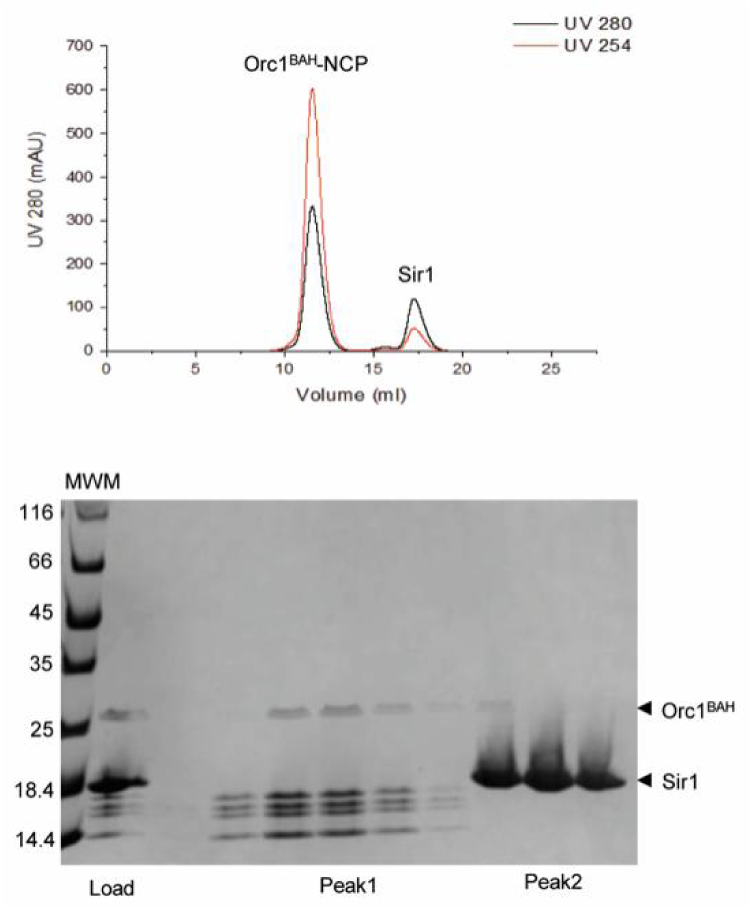
Sir1^OID^ cannot join the pre-formed Orc1^BAH^-NCP complex or compete for Orc1^BAH^ binding with NCP, as judged by an analysis using size-exclusion column chromatography. Top panel, UV absorbance chromatogram of the sizing column analysis; Bottom panel, SDS PAGE gel showing the content of the eluted fractions.

**Figure S2.**
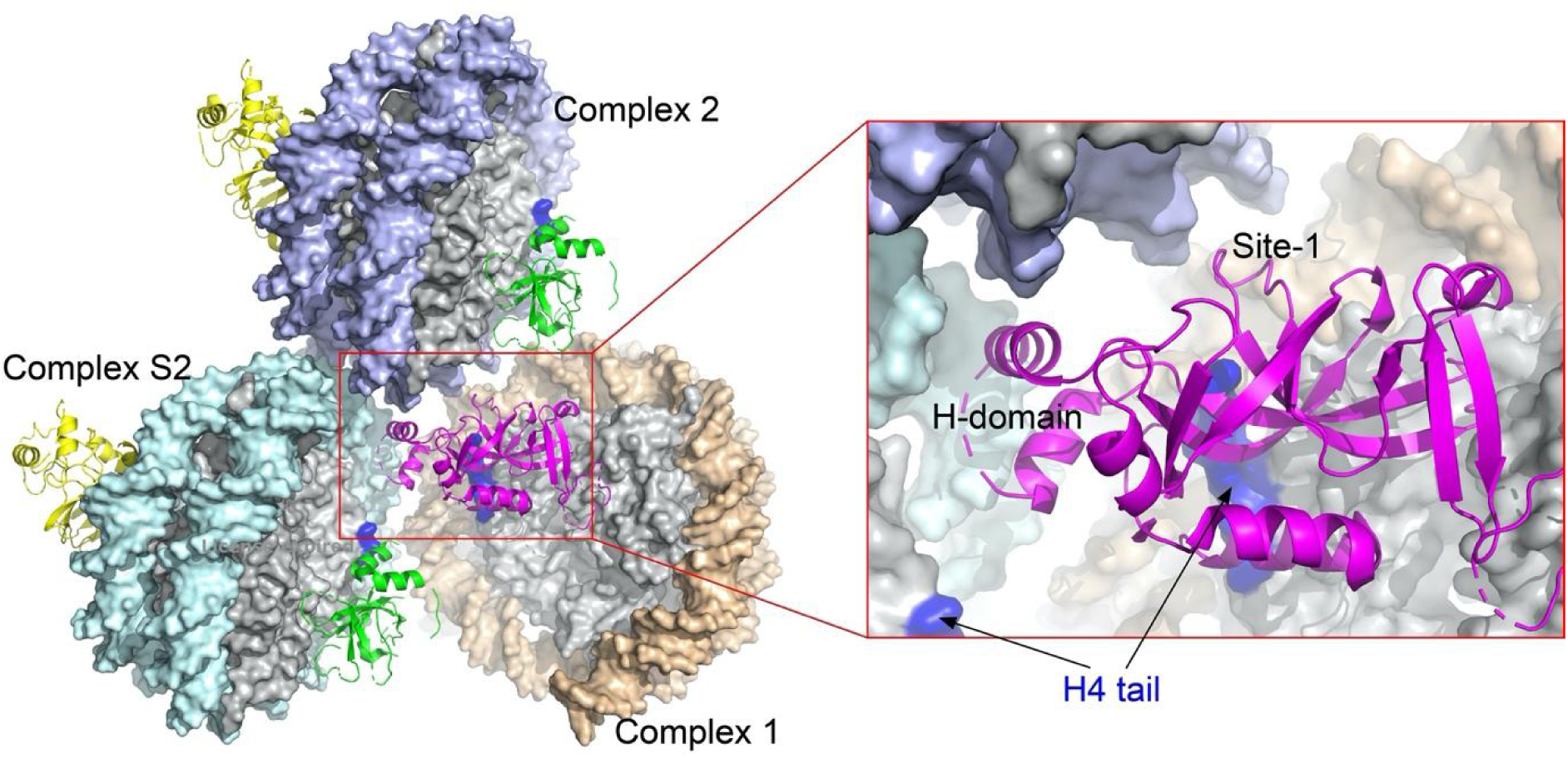
Sir1-binding region of Orc1^BAH^ engaged in crystal packing. Three closely packed Orc1^BAH^-NCP complexes are chosen to illustrate inter-complex interactions within and across the crystallographic asymmetric unit. The inset shows an enlarged view of the boxed area bordering the three complexes. The nucleosomal surface regions shaded blue indicate the location of N-terminal tails of histone H4. The Orc1^BAH^ molecule from the Orc1^BAH^-NCP complex 1 (colored magenta) contacts nucleosomal DNA from the symmetry-related Complex S2 (cyan) and that of the Complex 2 (light blue) in the same asymmetric unit through its H-domain. Its Site-1 loop is also near the Complex 2 DNA.

**Figure S3.**
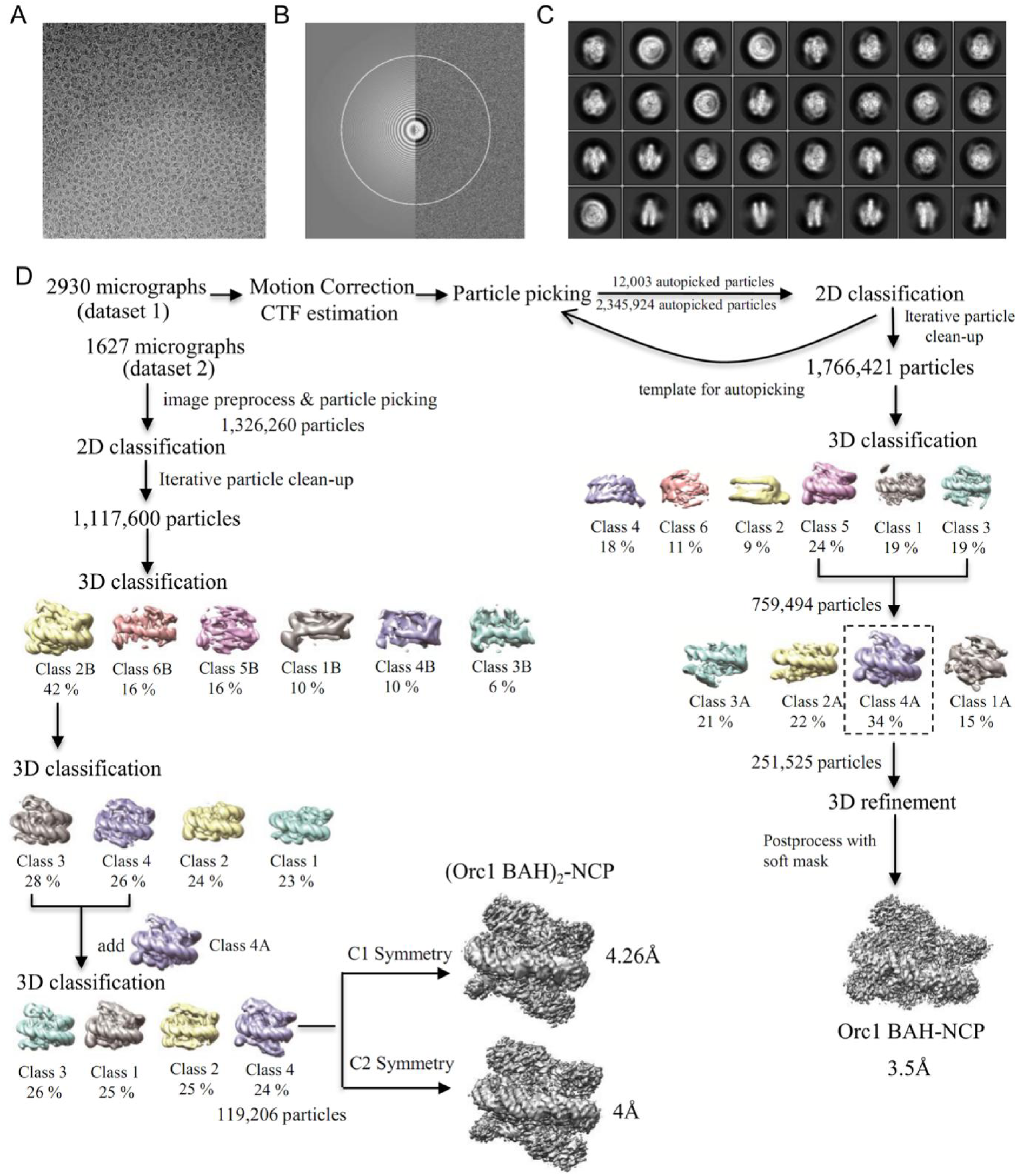
Workflow of cryo-EM data processing. (A) A representative electron micrograph low-pass filtered to 20 Å for improved particle visibility. (B) Contrast Transfer Function (CTF) estimation by the Gctf program. (C) Selected reference-free 2D class average. (D) Cryo-EM data processing procedure.

**Figure S4.**
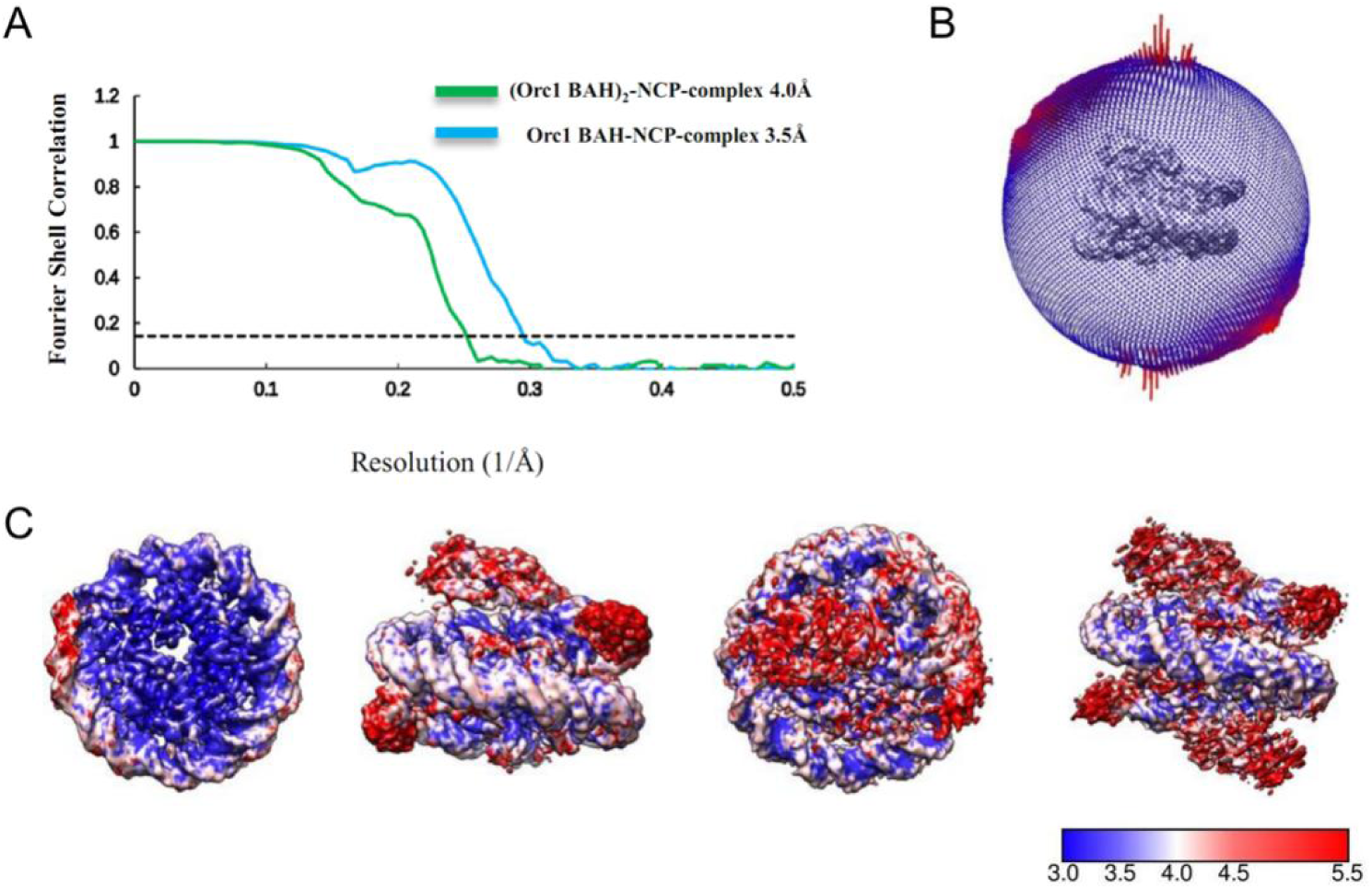
Cryo-EM map resolution. (A) Gold-standard Fourier correlation of the Orc1^BAH^-NCP complex and (Orc1^BAH^)_2_-NCP complex. (B) Angular distribution of cryo-EM particles in the final round of refinements of the Orc1^BAH^-NCP complex. (C) Local resolutions of the Orc1^BAH^-NCP (left two panels) and (Orc1^BAH^)_2_-NCP (right panels) complexes shown in bottom and side views, respectively. A heat bar coding resolution range is appended.

**Figure S5.**
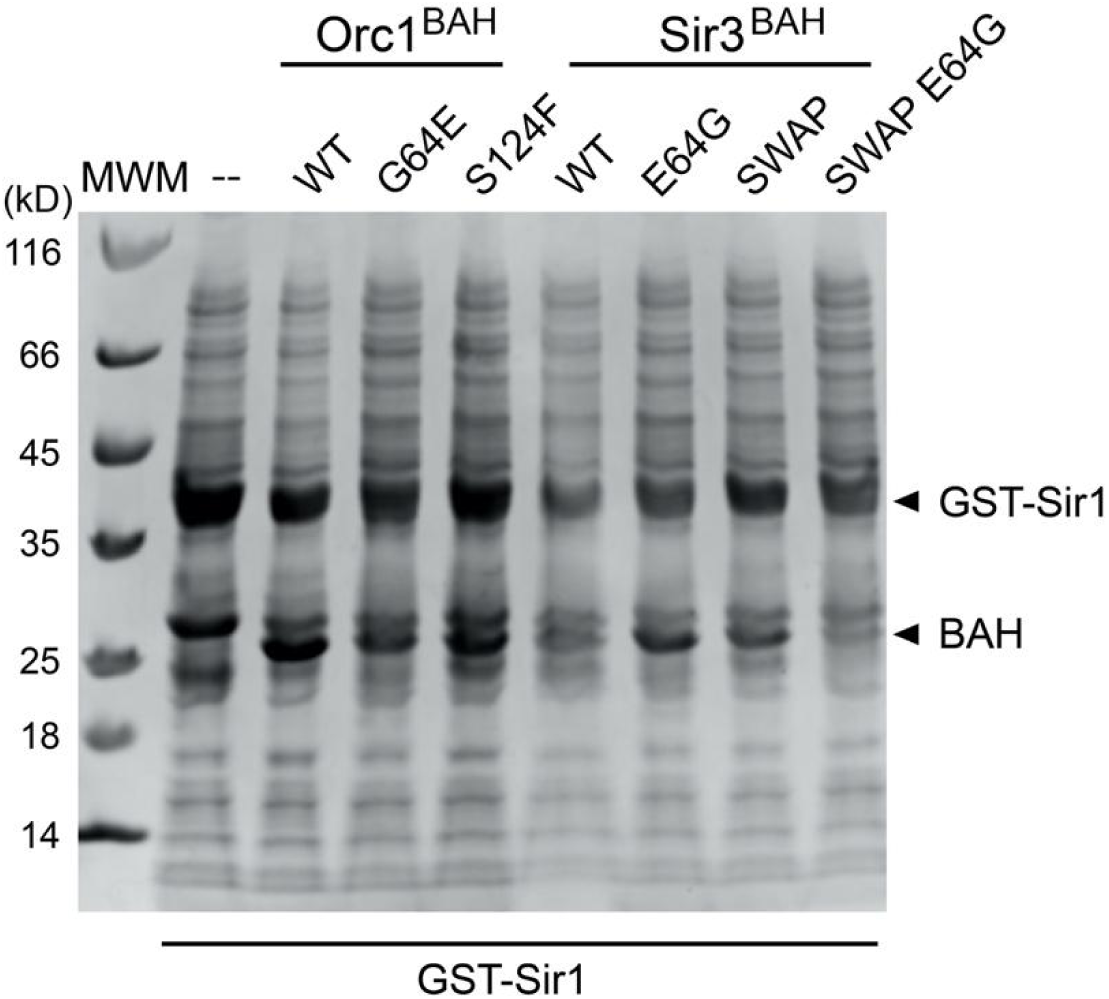
Inputs for GST-Sir1^OID^ pulldown of Orc1^BAH^ and Sir3^BAH^ variants used in Figure 3B. Cleared bacterial lysates of GST-Sir1^OID^ alone, or that co-expressed with the indicated wild-type or Orc1^BAH^ and Sir3^BAH^ mutants.

**Figure S6.**
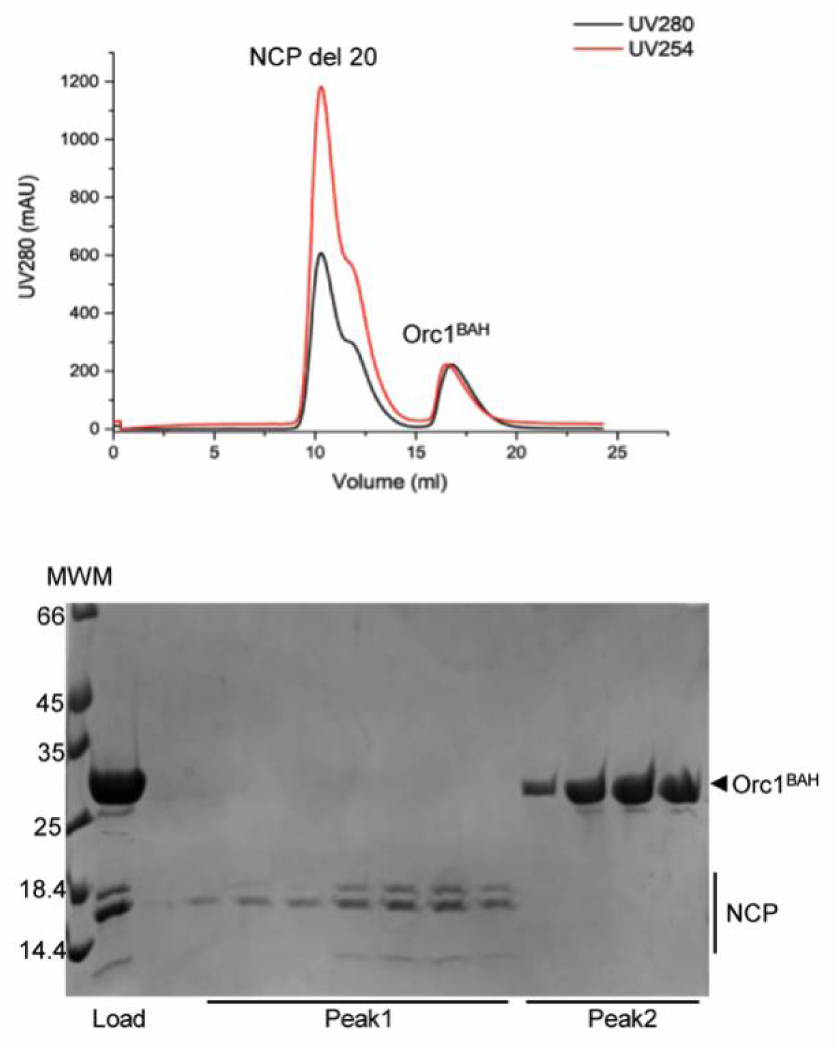
Orc1^BAH^ no longer binds NCP with deletion of 20 N-terminal residues of histone H4. Top panel, UV absorbance chromatogram of a size-exclusion column run of the mixture of Orc1^BAH^ and the mutant NCP; Bottom panel, SDS-PAGE gels shows that Orc1^BAH^ and the mutant NCP are eluted separately from the column.

**Table S1.**
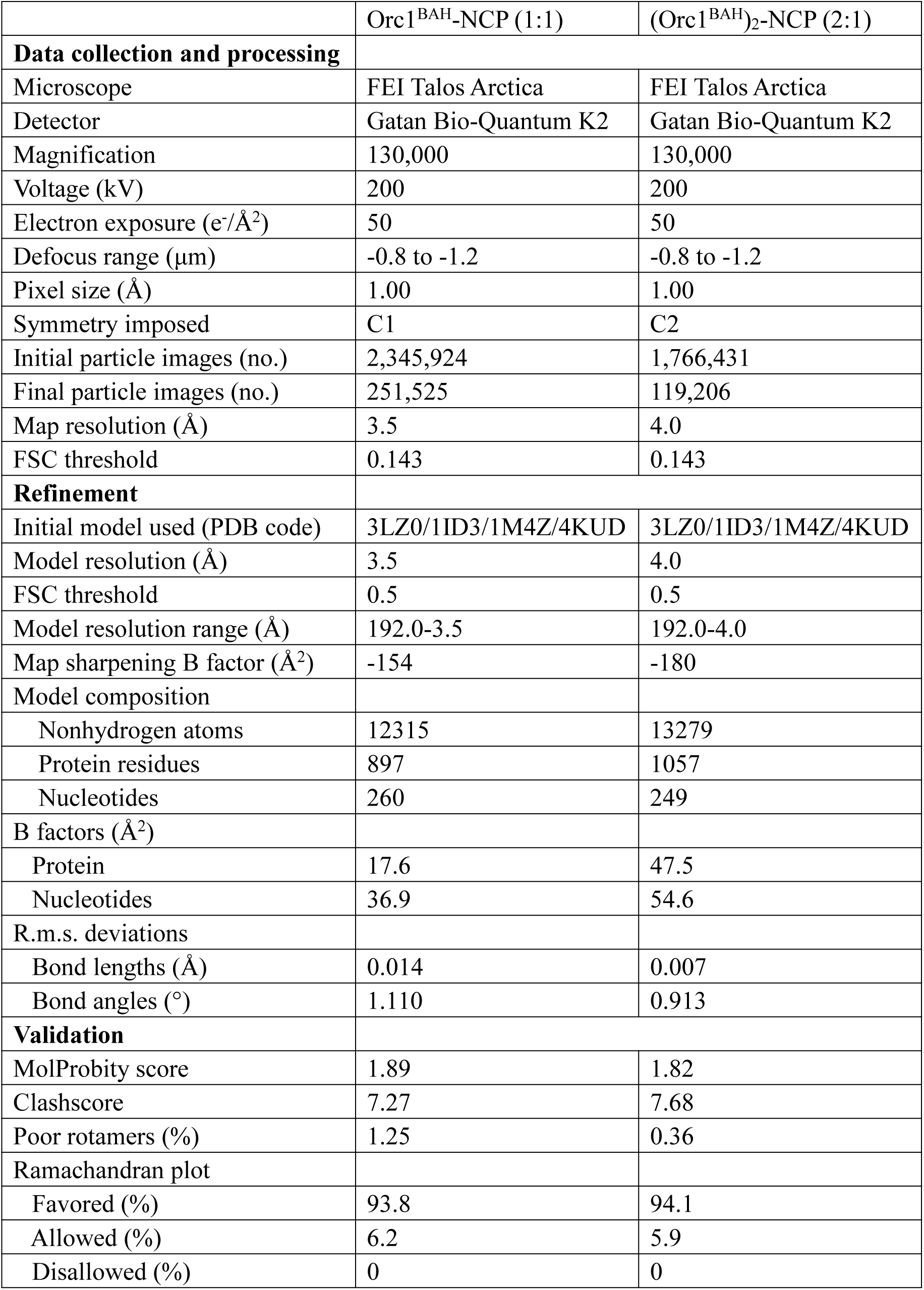
Cryo-EM data collection, refinement and validation statistics.

## Notes

### Competing Interest Statement

The authors have declared no competing interest.

