## Supplemental Table and Figures for "Nucleosome binding relinquishes the association of the BAH domain of Orc1 with Sir1"

**Table S1. Cryo-EM data collection, refinement and validation statistics**

| | $Orc1^{BAH}$-NCP (1:1) | $(Orc1^{BAH})_2$-NCP (2:1) |
|---|---|---|
| **Data collection and processing** | | |
| Microscope | FEI Talos Arctica | FEI Talos Arctica |
| Detector | Gatan Bio-Quantum K2 | Gatan Bio-Quantum K2 |
| Magnification | 130,000 | 130,000 |
| Voltage (kV) | 200 | 200 |
| Electron exposure ($e^-/Å^2$) | 50 | 50 |
| Defocus range (μm) | -0.8 to -1.2 | -0.8 to -1.2 |
| Pixel size (Å) | 1.00 | 1.00 |
| Symmetry imposed | C1 | C2 |
| Initial particle images (no.) | 2,345,924 | 1,766,431 |
| Final particle images (no.) | 251,525 | 119,206 |
| Map resolution (Å) | 3.5 | 4.0 |
| FSC threshold | 0.143 | 0.143 |
| **Refinement** | | |
| Initial model used (PDB code) | 3LZ0/1ID3/1M4Z/4KUD | 3LZ0/1ID3/1M4Z/4KUD |
| Model resolution (Å) | 3.5 | 4.0 |
| FSC threshold | 0.5 | 0.5 |
| Model resolution range (Å) | 192.0-3.5 | 192.0-4.0 |
| Map sharpening B factor ($Å^2$) | -154 | -180 |
| Model composition | | |
| Nonhydrogen atoms | 12315 | 13279 |
| Protein residues | 897 | 1057 |
| Nucleotides | 260 | 249 |
| B factors ($Å^2$) | | |
| Protein | 17.6 | 47.5 |
| Nucleotides | 36.9 | 54.6 |
| R.m.s. deviations | | |
| Bond lengths (Å) | 0.014 | 0.007 |
| Bond angles (°) | 1.110 | 0.913 |
| **Validation** | | |
| MolProbity score | 1.89 | 1.82 |
| Clashscore | 7.27 | 7.68 |
| Poor rotamers (%) | 1.25 | 0.36 |
| Ramachandran plot | | |
| Favored (%) | 93.8 | 94.1 |
| Allowed (%) | 6.2 | 5.9 |
| Disallowed (%) | 0 | 0 |

**Figure S1.**

Sir1$^{OID}$ cannot join the pre-formed Orc1$^{BAH}$-NCP complex or compete for Orc1$^{BAH}$ binding with NCP, as judged by an analysis using size-exclusion column chromatography. Top panel, UV absorbance chromatogram of the sizing column analysis; Bottom panel, SDS PAGE gel showing the content of the eluted fractions.

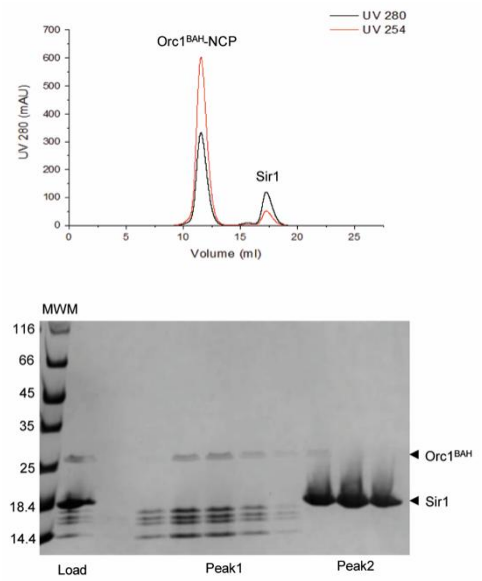


**Figure S2.**

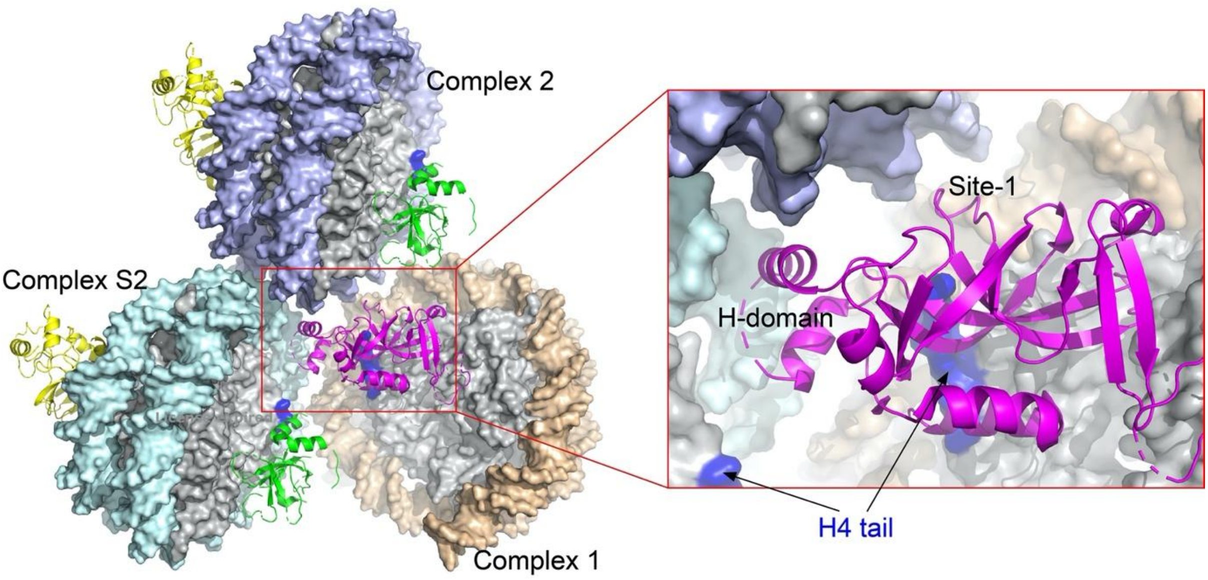


Sir1-binding region of Orc1$^{BAH}$ engaged in crystal packing. Three closely packed Orc1$^{BAH}$-NCP complexes are chosen to illustrate inter-complex interactions within and across the crystallographic asymmetric unit. The inset shows an enlarged view of the boxed area bordering the three complexes. The nucleosomal surface regions shaded blue indicate the location of N-terminal tails of histone H4. The Orc1$^{BAH}$ molecule from the Orc1$^{BAH}$-NCP complex 1 (colored magenta) contacts nucleosomal DNA from the symmetry-related Complex S2 (cyan) and that of the Complex 2 (light blue) in the same asymmetric unit through its H-domain. Its Site-1 loop is also near the Complex 2 DNA.

**Figure S3.**

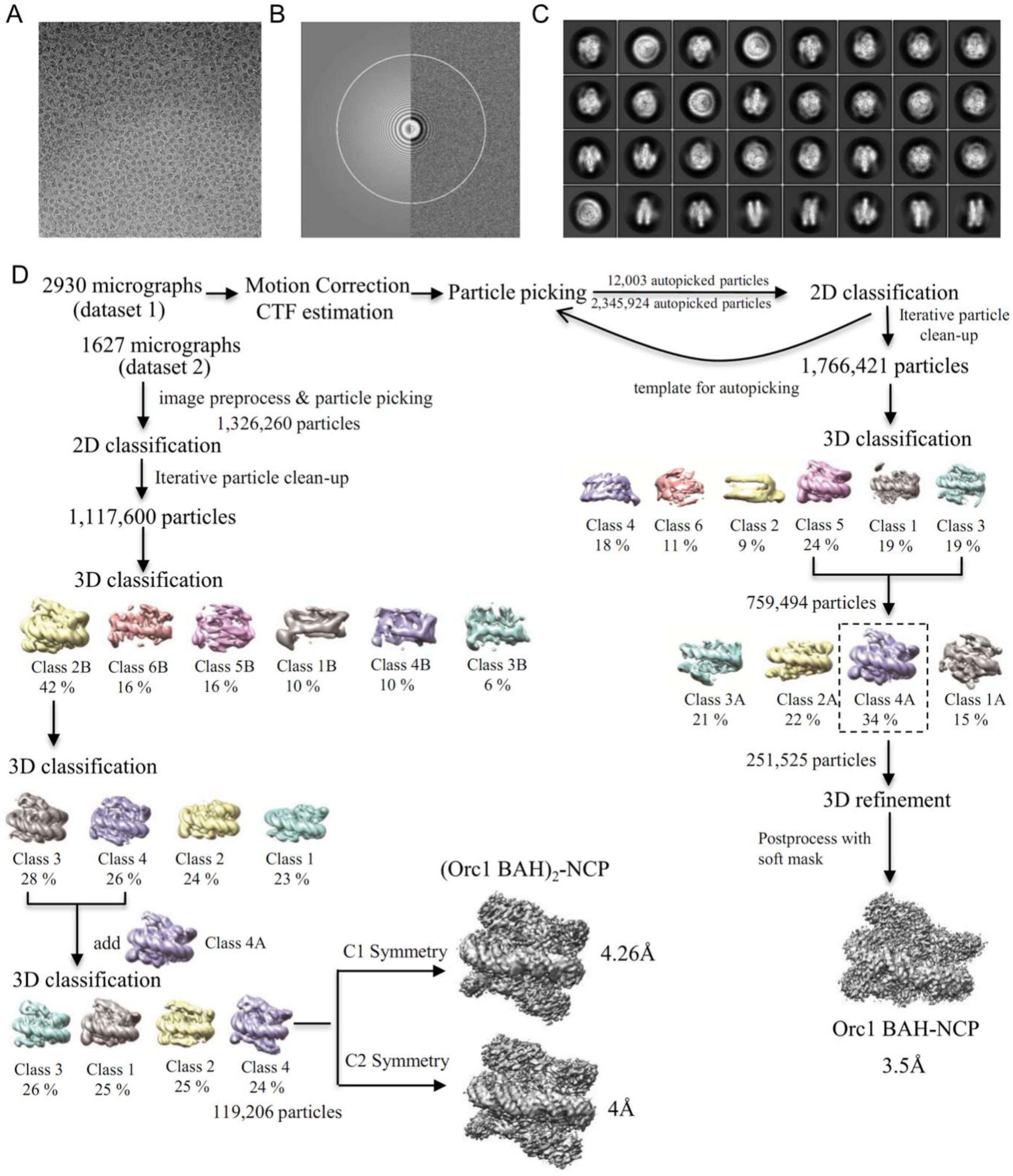


Workflow of cryo-EM data processing. (A) A representative electron micrograph low-pass filtered to 20 Å for improved particle visibility. (B) Contrast Transfer Function (CTF) estimation by the Gctf program. (C) Selected reference-free 2D class average. (D) Cryo-EM data processing procedure.

**Figure S4.**

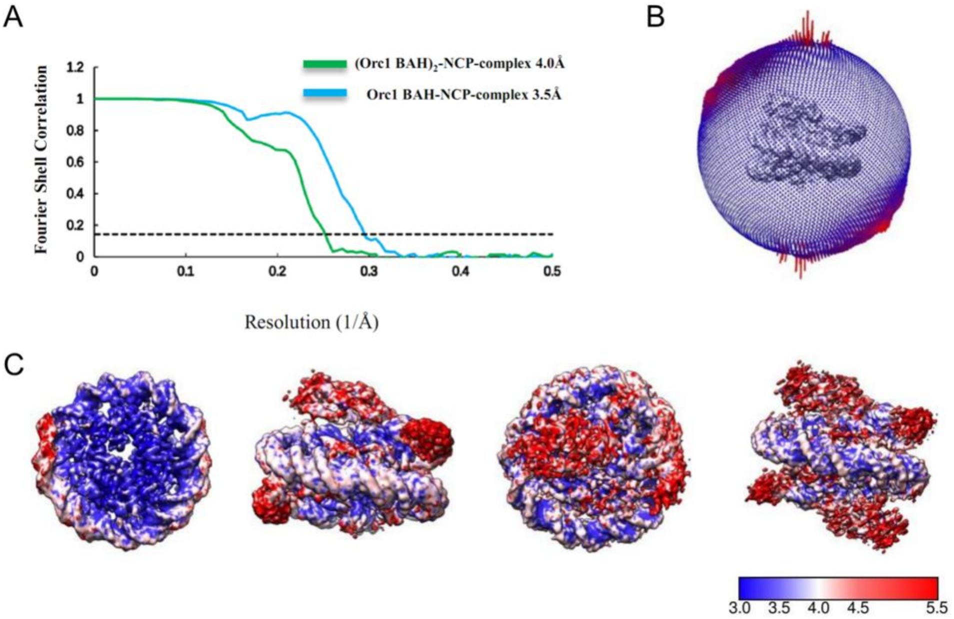


Cryo-EM map resolution. (A) Gold-standard Fourier correlation of the Orc1$^{BAH}$-NCP complex and (Orc1$^{BAH}$)$_2$-NCP complex. (B) Angular distribution of cryo-EM particles in the final round of refinements of the Orc1$^{BAH}$-NCP complex. (C) Local resolutions of the Orc1$^{BAH}$-NCP (left two panels) and (Orc1$^{BAH}$)$_2$-NCP (right panels) complexes shown in bottom and side views, respectively. A heat bar coding resolution range is appended.

**Figure S5.**

Inputs for GST-Sir1$^{OID}$ pulldown of Orc1$^{BAH}$ and Sir3$^{BAH}$ variants used in Figure 3B. Cleared bacterial lysates of GST-Sir1$^{OID}$ alone, or that co-expressed with the indicated wild-type or Orc1$^{BAH}$ and Sir3$^{BAH}$ mutants.

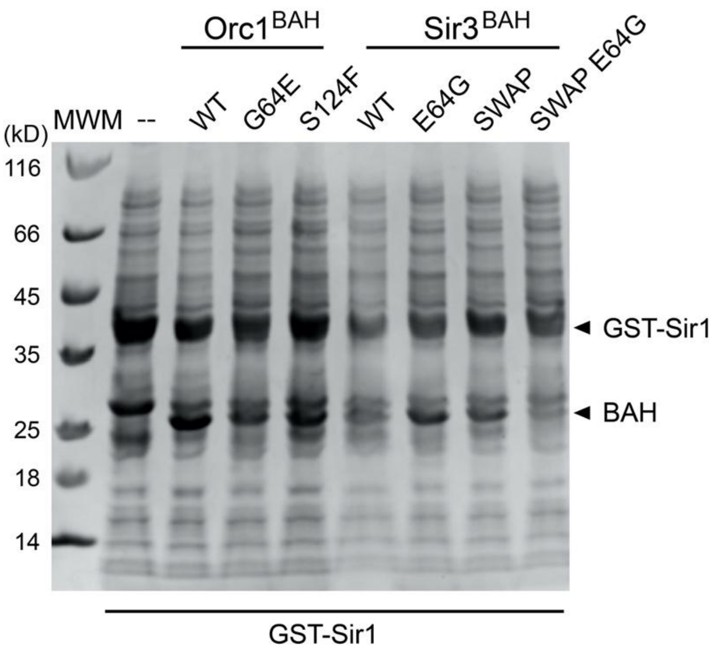


**Figure S6.**

Orc1$^{BAH}$ no longer binds NCP with deletion of 20 N-terminal residues of histone H4. Top panel, UV absorbance chromatogram of a size-exclusion column run of the mixture of Orc1$^{BAH}$ and the mutant NCP; Bottom panel, SDS-PAGE gels shows that Orc1$^{BAH}$ and the mutant NCP are eluted separately from the column.

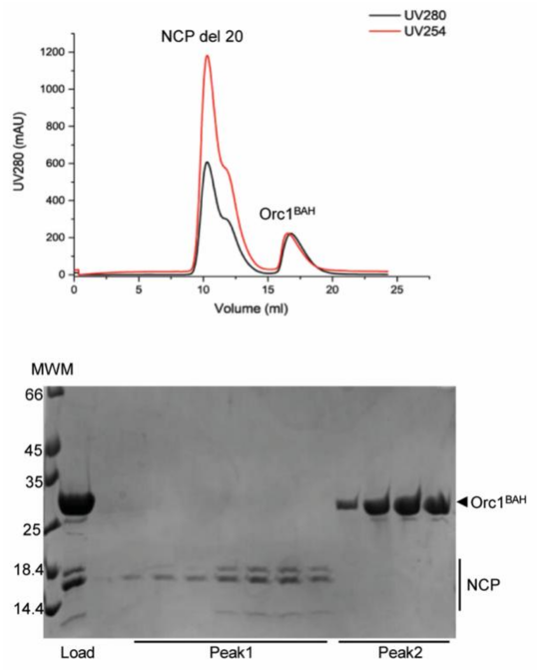
